# FADS1/2-mediated lipid metabolic reprogramming drives ferroptosis sensitivity in triple-negative breast cancer

**DOI:** 10.1101/2023.06.30.547227

**Authors:** Nicla Lorito, Angela Subbiani, Alfredo Smiriglia, Marina Bacci, Francesca Bonechi, Laura Tronci, Alessia Corrado, Dario Livio Longo, Marta Iozzo, Luigi Ippolito, Giuseppina Comito, Elisa Giannoni, Icro Meattini, Alexandra Avgustinova, Paola Chiarugi, Angela Bachi, Andrea Morandi

## Abstract

Triple-negative breast cancer (TNBC) has limited therapeutic options, is highly metastatic and characterized by early recurrence. Lipid metabolism is generally deregulated in TNBC and might reveal vulnerabilities to be targeted or used as biomarkers with clinical value.

Ferroptosis is a type of cell death caused by iron-dependent lipid peroxidation which is facilitated by the presence of polyunsaturated fatty acids (PUFA).

Here we identify fatty acid desaturases 1 and 2 (FADS1/2), which are responsible for PUFA biosynthesis, lipid susceptible to peroxidation, to be highly expressed in a subset of TNBC with a poorer prognosis. Lipidomic analysis, coupled with functional metabolic assays, showed that FADS1/2 high-expressing TNBC are susceptible to ferroptosis-inducing agents and that targeting FADS1/2 renders those tumors ferroptosis-resistant. These findings were validated *in vitro* and *in vivo* in mouse and human-derived clinically relevant models and in a retrospective cohort of TNBC patients.

**One sentence summary:** The availability of intracellular PUFA depends on FADS1/2 desaturases, expressed at higher levels in aggressive triple-negative breast cancers highly susceptible to ferroptosis.

## Introduction

Altered lipid metabolism is generally considered a common feature of breast cancers (Bacci et al., 2021). However, important differences have emerged in terms of lipid synthesis, upload, storage, and utilization between breast cancers with different biology, invasiveness, response to therapy, and prognosis (Koundouros & Poulogiannis, 2020). Intracellular lipids have a role as energetic substrates, structural components, or signaling molecules. A major component of lipids are fatty acids (FA) and their role within the cells is not only influenced by the total amount available but also by their length and complexity (i.e., presence of carbon-carbon double bonds). Indeed, the balance between unsaturated, mono (MUFA), and polyunsaturated FA (PUFA) is essential for cellular homeostasis and biological functions (Dyall et al., 2022). The insertion of double bonds is mediated by stearoyl-coenzyme A (stearoyl-CoA) desaturase 1 (SCD1), the key rate-limiting enzyme responsible for introducing the first double bond and therefore for MUFA formation. Subsequent double bonds are inserted by fatty acid desaturases 1 and 2 (FADS1/2) (Vriens et al., 2019; Xuan et al., 2022), which are thus key mediators of PUFA biosynthesis. Aberrant lipid desaturation is involved in tumor initiation and progression and can be effectively targeted to impair tumor growth, metastatic dissemination and relapse in preclinical models (Li et al., 2017; Peck et al., 2016; Ran et al., 2018). Particularly, PUFA have a major role in maintaining the architecture of the plasma membrane (Harayama & Shimizu, 2020). When in excess, PUFA are rapidly incorporated into the plasma membrane by long-chain acyl-CoA synthetase 4 (ACSL4) (Doll et al., 2017) and become vulnerable to lipid peroxidation. Peroxidized lipids alter the membrane bilayer, making it susceptible to rupture and causing ferroptosis, a unique form of iron-dependent cell death (Dixon et al., 2012). In addition, the FA composition of the membranes, particularly a higher presence of PUFA over that of MUFA, enhances ferroptosis sensitivity (Lei et al., 2022). Recently, many cell-autonomous pathways that prevent or revert lipid peroxidation have been shown to counteract ferroptosis induction. Among the mechanisms described, the antioxidant activity of the membrane-associated glutathione peroxidase 4 (GPX4) and the increased availability of the scavenger molecule glutathione (GSH) sustained by the SLC7A11-mediated import of cystine (Koppula et al., 2021), have a prominent role in regulating ferroptosis sensitivity.

Here we describe a cell-autonomous mechanism that exposes aggressive triple-negative breast cancer (TNBC) cells to ferroptosis susceptibility. We found that aggressive TNBC cells are characterized by higher expression of FADS1/2, leading to enhanced PUFA availability and thus becoming more sensitive to ferroptosis-promoting insults, a finding validated both *in vitro* and *in vivo*. Indeed, targeting FADS1/2 renders the aggressive TNBC cells resistant to ferroptosis. Finally, the addition of exogenous MUFA to the aggressive models exerts an anti-ferroptosis effect and prevents drug-induced cell death whereas increasing the availability of PUFA resensitizes the resilient models to ferroptosis execution.

## Materials and Methods

### Cell cultures and reagents

Human (BT20, MDA-MB-231) and murine (67NR, 4T07, 4T1, D2A1, D2A1-m1, D2A1-m2) female breast cancer cells were obtained from Prof. Clare Isacke (ICR, London, UK) and maintained at 37°C / 5% CO_2_ in phenol red Dulbecco’s Modified Eagle’s Medium (DMEM, Euroclone #ECB7501L) supplemented with 10% Fetal Bovine Serum (FBS, Euroclone #ECS0180L), 2 mmol/L glutaMAX (Gibco #35050-061), and 1% penicillin/streptomycin (Sigma #P4333). Cells were short tandem repeat tested, amplified, stocked, routinely subjected to mycoplasma testing, and once thawed were kept in culture for a maximum of 3-4 weeks. SB-204990 ((3R,5S)-rel-5-[6-(2,4-dichlorophenyl)-hexyl]-tetrahydro-3-hydroxy-2-oxo-3-furanacetic acid, #HY-16450), RSL3 ((1S,3R)-2-(2-chloroacetyl)-2,3,4,9-tetrahydro-1-[4-(methoxycarbonyl)phenyl]-1H-pyrido[3,4-b]indole-3-carboxylic acid, methyl ester, #HY-100218A), Erastin (2-[1-[4-[2-(4-chlorophenoxy)acetyl]-1-piperazinyl]ethyl]-3-(2-ethoxyphenyl)-4(3H)-quinazolinone, #HY-15763), A922500 (i.e., DGAT1 inhibitor, DGAT1i, (1R,2R)-2-[[4’-[[Phenylamino)carbonyl]amino] [1,1’-biphenyl]-4-yl]carbonyl]cyclopentanecarboxylic acid, #252801), PF-06424439 (i.e., DGAT2 inhibitor, DGAT2i, [(3R)-1-[2-[1-(4-chloro-1H-pyrazol-1-yl)cyclopropyl]-3H-imidazo[4,5-b]pyridin-5-yl]-3-piperidinyl]-1-pyrrolidinyl-methanone, monomethanesulfonate, #PZ0233), CP-24879 (4-(3-methylbutoxy)-benzenamine, monohydrochloride, #C9115), and ATGListatin N’-[4’-(dimethylamino)[1,1’-biphenyl]-3-yl]-N,N-dimethyl-urea, #HY-15859) were purchased from MedChemExpress, dissolved in DMSO and used at the indicated concentrations as described in the Figure Legends. TVB3166 (4-{1-[5-(4,5-dimethyl-2H-pyrazol-3-yl)-2,4-dimethyl-benzoyl]-azetidin-3-yl}-benzonitrile, #SML1694), SC-26196 (α,α-Diphenyl-4-[(3-pyridinylmethylene)amino]-1-piperazinepentanenitrile, #PZ0176), CAY10566 (2R-[[4’-[[(phenylamino)carbonyl]amino][1R,1’-biphenyl]-4-yl-carbonyl]-cyclopentanecarboxylic acid, #SML2980), etomoxir (2-[6-(4-chlorophenoxy)-hexyl]-oxirane-2-carboxylate, #E1905), ferrostatin-1 (fer-1, ethyl 3-amino-4-(cyclohexylamino)benzoate, #SML0583), deferoxamine mesylate salt (i.e., iron chelator, butanediamide, N’-[5-[[4-[[5-(acetylhydroxyamino)pentyl]amino]-1,4dioxobutyl]hydroxyamino]pentyl]-N-(5-aminopentyl)-N-hydroxy-, monomethanesulfonate, #D9533), rosiglitazione (i.e., ACSL4 inhibitor, ACSL4i, 5-[[4-[2-(Methyl-2-pyridinylamino)ethoxy]phenyl]methyl]-2,4-thiazolidinedione, #R2408) from Sigma-Aldrich were also dissolved in DMSO. TOFA (5-(tetradecyloxy)-2-furoic acid, #sc200653) was purchased from Santa Cruz Biotechnology and dissolved in DMSO.

The exogenously administrated FA adrenic acid (C22:4; all-cis-7,10,13,16-docosatetraenoic acid, #D3659), erucic acid (C22:1; cis-13-docosenoic acid, #E3385), and palmitoleic acid (C16:1; *cis*-9-hexadecenoic acid, #P9417) were purchased from Sigma-Aldrich and dissolved in ethanol.

### Mouse models and *in vivo* experiments

6-weeks-old female BALB/c mice were orthotopically inoculated with 4T1 (2.5 × 10^5^ cells, n=6) resuspended in 50 μL of Dulbecco’s phosphate buffered saline (PBS, Euroclone, #ECB4053) in a single flank injection. Tumors were grown to ∼10 mm diameter (day 8) and then the mice were randomly assigned to the vehicle or to RSL3-treated group. RSL3 (40 mg/kg) was injected intraperitoneally every other day in 50 μL of a mixture of 40% PEG300, 45% saline, 10% DMSO, and 5% Tween-80 starting from day 8. Tumor volume was monitored from day 7 onward by caliper measurements of the two largest tumor diameters. Volumes were calculated according to the formula: a×b2×π/6, where a and b are orthogonal tumor diameters. Animal weight was measured three times a week. Animals were culled at day 23 when tumor volumes of the vehicle treated group reached the maximum tumor size allowed by the Animal Welfare license (License number 741/2022).

### Survival assay

Human and murine breast cancer cells were seeded into 12-well plates at 30-50.000 cells/well in either standard conditions (see cell cultures and reagents) or experimental conditions such as RNA knockdown (i.e., siFADS1) or drug administration (e.g., 0.1 μM of the GPX4 inhibitor, RSL3), as described in the Figures and relative legends. Unless specified otherwise, 24 hours post-procedure cells were extensively washed with PBS, fixed with 4% formaldehyde, and subjected to the colorimetric evaluation of cell proliferation and viability using crystal violet (triphenylmethane dye 4-[(4-dimethylaminophenyl)-phenyl-methyl]-*N*,*N*-dimethyl-alanine, Sigma-Aldrich #548-62-9), thus dried overnight (ON). The incorporated amount of crystal violet within the adherent living cells was solubilized with a maximum of 500 μl/well of 2% SDS and quantified by measuring the absorbance at 595 nm using a microplate reader. Alternatively, cell counts were performed in triplicate by three analysts under a 10x objective according to the standard methodology.

### RNAi transfection

4T1 cells were seeded into 6-well plates (3 × 10^5^ per well) to achieve 70% confluence the following day, when cells were transfected with either 100 nmol/L siRNA targeting FADS1 (siFADS1, GE Healthcare Dharmacon), or the respective negative controls (non-targeting small interfering RNA, siCTR, GE Healthcare Dharmacon) using Lipofectamine RNAiMAX Reagent (Thermo Fisher Scientific #13778-150) and Opti-MEM (GIBCO #31985062) accordingly to manufacturer’s instructions. The functional analyses were performed 24 hours after transfection as described in the Figure Legends.

### Western blotting analysis

Human and murine breast cancer cells were washed with PBS and lysed on ice using 1x Laemmli Sample Buffer (Biorad #161-0737) supplemented with protease (Sigma-Aldrich #P8340) and phosphatase inhibitors (#P0044). Protein concentrations were measured using BCA assay (Sigma-Aldrich #1003290033). 40–50 μg of cell lysate were loaded in precast SDS-PAGE (sodium dodecyl sulfate–polyacrylamide gel electrophoresis) gels (Biorad #456-8096) and then transferred onto nitrocellulose membrane by Trans-Blot Turbo Transfer Pack (Biorad #170-4157). The immunoblots were incubated in PBS-T (PBS with 0.05% tween-20) containing 2% non-fat dry milk at room temperature for 1 hour and then probed with primary and appropriate secondary antibodies. The following antibodies were used: ACLY (Santa Cruz Biotechnology #sc-517267), FASN (Santa Cruz Biotechnology, #sc-55580), PLIN2 (Cell Signaling Technology #45535), FADS1 (Sigma-Aldrich, #SAB2100744), FADS2 (Sigma-Aldrich #SAB1303849), SCD1 (Santa Cruz Biotechnology, #sc-515844), SCD2 (Santa Cruz Biotechnology, #sc-518034) ACSL4 (Santa Cruz Biotechnology, #sc-365230), HSP90 (Santa Cruz Biotechnology #sc-69703), and β-actin (Cell Signaling #3700), all diluted 1:1000 in PBS-T containing 2% non-fat dry milk.

### RNA extraction and Quantitative Real Time PCR (qRT-PCR) analysis

Total RNA was extracted using RNeasy plus kit (QIAGEN #74134), quantified using Nanodrop 1000 (Thermo Fisher Scientific), and 500 ng were reverse transcribed using the iScript gDNA Clear cDNA Synthesis Kit (Biorad #172-5035). qRT-PCR was performed using the CFX96 Touch Real-Time PCR Detection System (Biorad) using TaqMan Universal PCR Master Mix (Thermo Fisher Scientific #4305719). The following probes were used: ACLY, FASN, PLIN2, and ACSL4 (Thermo Fisher Scientific). Data were normalized on TBP (TATA-Box Binding Protein) or GAPDH (Thermo Fisher Scientific). The relative quantity was determined using ΔΔCt by the CFX Maestro software (BioRad).

### Radioactive assay

Breast cancer cells (8 ×10^4^ cells/well) were seeded into 12-well plates. To analyze the incorporation of ^14^C-glucose into lipids, culture media were supplemented ON with 1 mCi ^14^C-glucose (Perkin Elmer #NEC042V250UC). Cells were then washed 3 times in ice cold PBS and lysed in RIPA buffer (Thermo Fisher Scientific #89900). Samples were first resuspended in 4 volumes of a CHCl3:MeOH (1:1) solution and then an additional volume of dH2O was added. The solution was then centrifuged at 1000 rpm for 5-10 minutes at room temperature. The lower phase (i.e., lipids) was collected, transferred to a scintillation vial, and the incorporated radioactive glucose derived signal was measured on the scintillation counter and normalized to the total protein content.

### Seahorse-based oxidative phenotyping

67NR, 4T07, and 4T1 cells were seeded in XFe96 cell culture plates with 3 × 10^4^ cells per well (10 technical replicates) in 80 μL of culture medium, subjected to the experimental conditions described in the Figures and incubated at 37°C. 24 hours post seeding, the culture medium was replaced with 180μL of XF DMEM supplemented with 25 mM glucose (Sigma #G8644) and 2 mM glutaMAX. Cells were incubated for 1 hour at 37°C in a non-CO_2_ incubator to allow to pre-equilibrate with the XF DMEM. The oxygen consumption rate (OCR) was quantified using the Seahorse Extracellular Flux Analyzer (XFe96, Agilent Technologies). An accurate titration with the uncoupler FCCP was performed for each cell type. The addition of the ATP synthase inhibitor oligomycin (1.5 μM), the proton uncoupler FCCP (1 μM), the respiratory complex I inhibitor rotenone (0.5 μM), and the respiratory complex III inhibitor antimycin A (0.5 μM) was carried out at the times indicated. This assay measures the cellular substrate oxidation by assessing changes in the OCR of live cells when the entry of specific mitochondria (i.e., TCA cycle) fueling substrates is impaired. Indeed, the XF Cell Mito Stress Test was combined with substrate pathway specific inhibitors: etomoxir (4 μM) for long-chain fatty acids (LCFA) through inhibition of carnitine palmitoyl transferase 1a (CPT1a), UK5099 (2 μM) for glucose and/or pyruvate through inhibition of the mitochondrial pyruvate carrier (MPC), and BPTES (3 μM) for inhibition of glutamine through glutaminase 1 (GLS-1). Protein quantification was used to normalize the results. Basal respiration is calculated as the last rate measurement before oligomycin injection – non-mitochondrial respiration rate. Maximal respiration is calculated as the maximum rate measurement after FCCP injection – non-mitochondrial respiration rate.

### Oroboros O2k-FluoRespirometer

Oxygen consumption was analyzed in 2 mL glass chambers at 37°C using the Oroboros Oxygraph-2K high-resolution respirometer (Oroboros Instruments, Innsbruck, Austria) and the substrate, uncoupler, inhibitor, titration (SUIT) protocol (D009) (Ye & Hoppel, 2013). The oxygen flux normalized on the cell number is calculated as the negative time derivative of the oxygen concentration, measured in sealed chambers, and normalized on the instrumental background (measured in a dedicated experiment before assaying the cells). Murine breast cancer cells cultured in basal condition and treated with 40 µM Etomoxir or vehicle (DMSO) for 30 minutes were subjected to respirometry analysis. After instrumental air calibration, 5-7 x10^5^ cells resuspended in complete culture medium were introduced into the chambers and the basal respiratory activity was measured as routine respiration (R). The LEAK state (L) represents the non-phosphorylating state of uncoupled respiration due to proton leak, proton and electron slip, and cation cycling (Pesta & Gnaiger, 2012) after the inhibition of ATP synthase by oligomycin administration (5 nM). The capability of the electron transfer system (ETS) was measured by uncoupler titrations using the uncoupler Carbonyl Cyanide 3-ChloroPhenylhydrazone (CCCP; 1.5 µM/titration steps) as the readout of the maximal capacity of oxygen utilization (E). The residual oxygen consumption (ROX) that remains after the inhibition of ETS was determined by antimycin A injection (2.5 µM). Data acquisition and analysis were performed using DatLab software (Oroboros Instrument, Innsbruck, Austria) and the oxygen fluxes recorded in the individual titration steps were corrected for ROX.

### Lipidomic analysis

Lipids were extracted using a single-step extraction protocol with methanol and chloroform as described previously (Cattaneo et al., 2021). Briefly, cell pellets (equivalent to 1–3 × 10^3^ cells) were resuspended in 200 µL of MilliQ water and mechanically disrupted by passing 20 times through a 26 G syringe needle. Proteins were extracted from 20 μL lysate by adding 5 μL of lysis buffer (10% NP40, 2% SDS in PBS) and quantified by BCA protein assay kit. Lipids were extracted starting from equivalent lysate corresponding to 50 μg of proteins. Lysates were made up to 170 μL with cold water and spiked in with 1 μL of SPLASH^®^ LIPIDOMIX^®^ Mass Spec Standard. Lipid extraction was performed by adding 700 μL of methanol and subsequently 350 μL of chloroform. Samples were mixed on the orbital shaker for 15 minutes at 4 °C. After that, 350 μL of water/chloroform (1:1 v/v) were added to each suspension and centrifuged at 10,000× g for 10 minutes at 4 °C. The organic phase was recovered, dried out and finally resuspended in 25 μL of a buffer composed by 90% of Lipidomics Buffer A (95% of mobile phase A = ACN:H_2_O 40:60; 5 mM NH_4_COOCH_3_; 0.1% HCOOH and 5% of mobile phase B = Isopropanol:H2O 90:10; 5mM NH_4_COOCH_3_; 0.1% HCOOH) and 10% of ethanol. 1 μL was injected on the nLC Ekspert LC400 set in nano configuration coupled with the mass spectrometer Triple TOF 6600 (AB Sciex). Chromatography was performed using an in-house packed nanocolumn Kinetex EVO C18, 1.7 μm, 100 A (Phenomenex, Torrance, CA, USA), 0.75 × 100 mm at room temperature. The gradient started from 5% of mobile phase B and was linearly increased to 100% B in 5 minutes, maintained for 45 minutes, then returned to the initial ratio in 2 minutes and maintained for 8 minutes at a flow rate of 150 nL/min. Samples were analyzed in positive mode with electrospray ionization. Spectra were acquired by full-mass scan from 200 to 1700 m/z and information-dependent acquisition (IDA) from 50 to 1800 m/z (top 8 spectra per cycle). The de-clustering potential was fixed at 80 eV, and collision energy at 40 eV, target ions were excluded for 20 s after 2 occurrences. Wiff files were processed using the open-source software MS-DIAL version 4.8 with manual inspection of peak integrations. Peak areas were normalized by using the sum of lipid species areas for each sample. For lipid unsaturation analysis, normalized lipid areas were grouped into lipid classes and further sub-grouped according to the number of double bonds. For each subgroup, averages were used to calculate the PUFA/MUFA ratio.

### Confocal image acquisition and analysis

Breast cancer cells were seeded onto glass coverslips (2-3 × 10^5^ per well of a 6-well plate) to have a 50% of confluence. The day after manipulation cells were stained at 37°C (i) for 15 minutes with BODIPY^493/503^ (Thermo Fisher Scientific #D3922) to reveal LD content, or (ii) 1 hour with BODIPY^581/591^-C11 to evaluate lipid peroxidation (Thermo Fisher Scientific #D3861) and then fixed with 4% formaldehyde for 15 minutes. For nuclei staining, fixed cells were incubated with DAPI (Thermo Fisher Scientific #D3571) for 10 minutes at room temperature. Sample images were acquired using TCS SP8 microscope (Leica Microsystems) with LAS-AF image acquisition software. The quantification of LD was performed using CellProfiler software.

### Flow cytometry analysis

Breast cancer cells (3-5 ×10^4^ cells/well) were seeded into 12-well plates and subjected to the experimental procedure. The day after, cells were stained at 37°C for (i) 15 minutes with BODIPY^493/503^ to reveal LD content, or (ii) 1 hour with BODIPY^581/591^-C11 to evaluate lipid peroxidation. Live cells resuspended in PBS with 0.1% FBS were subjected to flow cytometry analysis using a FACSCanto II (BD Biosciences). 1×10^4^ cells were analyzed for the median of fluorescence intensity (MFI) of specific probe.

### Reactive Oxygen Species (ROS) analysis

Murine breast cancer cells (3-5×10^4^ cells/well) were seeded into 12-well plates and subjected to the experimental conditions described in Figures. Following a 24-hour incubation, cells were stained with CellROX (Thermo Fisher Scientific #C10444), DCFDA (Sigma-Aldrich #D6883) or MitoSOX (Thermo Fisher Scientific #M36008) to evaluate ROS cytoplasmatic and mitochondrial content and incubated at 37°C in the dark for 30 minutes. Then, cells were lysed with RIPA buffer and fluorescence was measured on a microplate reader at 485 nm excitation (Ex) and 520 nm emission (Em) for cellROX, at 485/535 nm Ex/Em for DCFDA, at 510/580 nm Ex/Em for MitoSOX.

### GSH/GSSG measurement

Murine breast cancer cells were seeded in 12-well plates (3-5×10^4^ cells/well) and GSH and GSSG concentration were measured after 2 hours of RSL3 1μM incubation according to manufacturer’s instructions (GSH/GSSG-Glo Assay, Promega #V6611). Luminescence was read using a luminometer and was proportional to the amount of GSH. A twofold adjustment was required for GSSG concentration because each mole of oxidized GSSG upon reduction produces two moles of GSH in this assay. GSH/GSSG ratio was calculated directly from luminescence measurements (in relative light units, RLU).

### *In silico* analysis - Analysis of human datasets

For *FADS1*, *FADS2* and *FADS1*+*FADS2* survival analyses, the curated dataset of ER-, PR- (assessed by immunohistochemical staining), HER2- (assessed by array) breast cancers was created using Km-plotter (http://kmplot.com) and included the overall survival (OS) data of patients belonging to the following datasets: GSE81538 (Brueffer et al., 2018), GSE96058 (Brueffer et al., 2018). For *FADS1* (208964_s_at), *FADS2* (202218_s_at) or both survival analysis, the curated dataset included the relapse free survival (RFS) data of patients belonging to the following datasets: GSE21653 (Sabatier et al., 2011), GSE31519 (Rody et al., 2011), GSE25066 (Hatzis et al., 2011), GSE2603 (Amaro et al., 2016), GSE69031 (Chin et al., 2006), GSE19615 (Li et al., 2010), E-MTAB-365 (Guedj et al., 2012), GSE65194 (Maubant et al., 2015). For FADS1+FADS2 survival analysis, the weighted (1:1) mean expression of the selected genes was used and patients were dichotomized into high and low expressing using the best cutoff threshold (Lánczky & Győrffy, 2021).

### Statistics and reproducibility

Statistics were performed using Prism 9 (GraphPad Software). Lipidomics statistical analysis, heatmaps and volcano plots were made with Metaboanalyst 5.0 (https://www.metaboanalyst.ca/). Unless stated otherwise, all numerical data are expressed as the mean ± standard error of the mean (SEM). All experiments were conducted at least 3 times independently, with 3 or more technical replicates for each experimental condition tested. Unless stated otherwise, comparisons between 2 groups were made using the two-tailed, unpaired Student’s t-test. Comparisons between multiple groups were made using one-way or two-way analysis of variance (ANOVA). Bonferroni and Dunnett post-testing analysis with a confidence interval of 95% was used for individual comparisons as reported in figure legends. Multivariate Cox analyses on the cohort of patients analyzed were generated using KM-plotter. Statistical significance was defined as: * *P*<0.05; \*\**P*<0.01; \*\*\**P*<0.001; \*\*\*\**P*<0.0001; when differences were not statistically significant or the comparison not biologically relevant, no indication was reported in the figures.

## Results

### Altered lipid metabolism is a feature of more aggressive TNBC cells

We used isogenic 67NR, 4T07 and 4T1 TNBC cells, which were previously derived from the same primary tumor and characterized by diverse tumorigenic and metastatic capacities (Avgustinova et al., 2016) (**Figure 1a**). Specifically, when injected orthotopically, 67NR cells do not metastasize, whereas 4T07 tumors seed micrometastases only in the lung, while 4T1 are overtly metastatic TNBC cells (Avgustinova et al., 2016). It has been previously reported that 4T1 cells are characterized by high metabolic flexibility, a trait that may correlate with their increasing aggressive ability (Simões et al., 2015).

**Figure 1.**
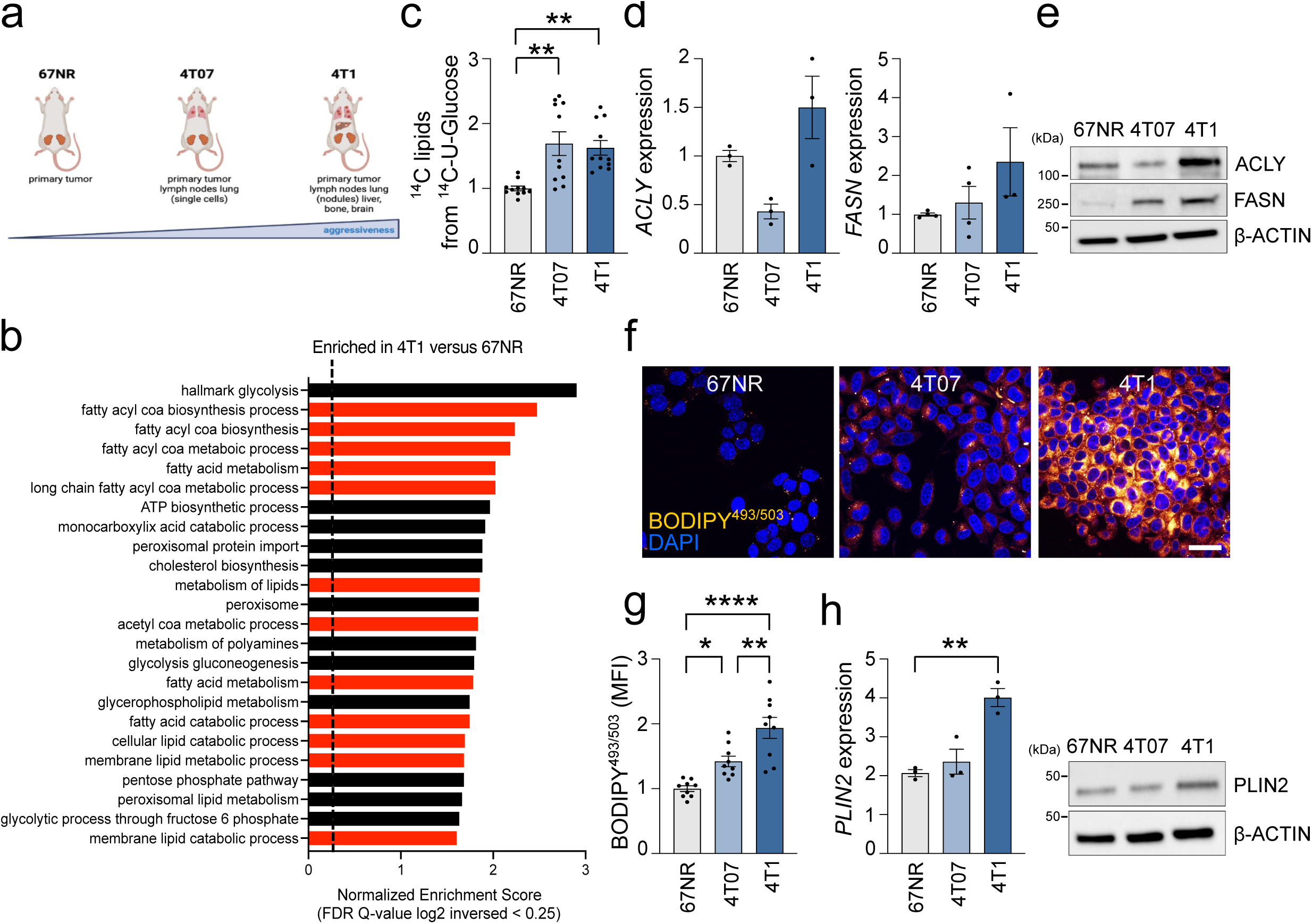
Altered lipid metabolism is a feature of more aggressive TNBC cells. (a) Schematic representation of the metastatic potential of the 4T1 cell line series. All the cell lines of the 4T1 series have been orthotopically injected into wild-type BALB/c mice, showing different aggressiveness and metastatic potential as here represented. (b) Gene Set Enrichment analysis (GSEA) results of the top gene sets showing a positive association with the 4T1 versus 67NR gene expression profile (GSE236033). NES, normalized enrichment score. FDR, false discovery rate. (c) 67NR, 4T07, and 4T1 breast cancer cells were cultured overnight (ON) in a medium containing ^14^C-U-(uniformly) radioactive labeled glucose. Lipids were extracted and the radioactive signal was measured to monitor the amount of ^14^C-U-glucose that is incorporated into lipids, as described in Methods. Each value was normalized on protein content. Data represent means ± standard errors of the means (SEMs), n = 3. One-way ANOVA, Dunnett corrected. (d) Murine breast cancer cells were analyzed by quantitative real-time polymerase chain reaction (qRT-PCR) analysis using the assays described in the figure. Fold relative enrichment is shown using the non-metastatic cells as comparator. Data represent means ± SEMs. One-way ANOVA, Dunnett corrected. (e) Total protein lysates from less to more aggressive cells were subjected to western blot (WB) analysis with the antibodies indicated. (f) TNBC cells were subjected to confocal analysis. Representative pictures of BODIPY^493/503^ stained cells are shown (orange/yellow: LD; blue: DAPI, nuclei). Scale bar, 10 μm. Data represent means ± SEMs. One-way ANOVA, Dunnett corrected. (g) The 4T1 series cells were subjected to cytofluorimetric analysis. FACS analysis of the mean fluorescence intensity (MFI) of the populations positive for BODIPY^493/503^ was reported. Data represent means ± SEMs. One-way ANOVA, Dunnett corrected. (h) Murine breast cancer cells were analyzed by qRT-PCR and WB analysis using the assay and the antibodies described in the figure. Fold relative enrichment is shown using the non-metastatic cells as comparator. Data represent means ± SEMs. One-way ANOVA, Dunnett corrected. Each dot represents a biological replicate. * *P*<0.05; ** *P*<0.01; **** *P*<0.0001.

To identify whether the observed metabolic features were stochastic or rather supported by a general transcriptomic and metabolomic reprogramming, whole genome expression analysis was performed on tumor cells FACS isolated from 4T1 and 67NR tumors. Subsequent Gene Set Enrichment analysis (GSEA) confirmed that the HALLMARK_GLYCOLYSIS gene set (MSigDB M5937) was positively correlated (NES=2.9; FDR=0.001) with the 4T1 transcriptome and surprisingly revealed a positive correlation with several gene sets related to fatty acid (FA) biosynthesis and lipid metabolism, indicating that glucose can also contribute to citrate production in the TCA cycle, thus possibly promoting *de novo* lipid production and utilization (**Figure 1b**).

Therefore, to functionally assess the cells’ ability to foster lipid biogenesis, we pulsed 67NR, 4T07, and 4T1 cells with ^14^C-glucose before recovering the lipids fraction. The more aggressive cell lines showed an increase in glucose-dependent lipid biogenesis when compared to 67NR (**Figure 1c**). Accordingly, qRT-PCR and western blot analyses revealed increased expression levels of the key lipogenic enzymes ATP Citrate Lyase (ACLY) in 4T1 cells and Fatty Acid Synthase (FASN) in both 4T07 and 4T1 cells when compared to 67NR cells (**Figure 1d,e**). This enhanced ability to sustain lipogenesis was paralleled by an increased lipid accumulation in form of lipid droplets (LD) content in the more aggressive cells as visualized by confocal microscopy (**Figure 1f**) and flow cytometry (**Figure 1g**) analyses using the fluorescent neutral lipid dye BODIPY^493/503^. LD increased accumulation was also supported by increased levels of mRNA and protein expression of Perilipin 2 (PLIN2), a key protein responsible for LD’ stability and formation (**Figure 1h**). To exclude that the observed metabolic phenotype was cell line specific, we validated the findings in two isogeneic spontaneously metastatic sublines, D2A1-m1 and D2A1-m2, generated from the poorly metastasizing murine-derived D2A1 cell line by serial *in vivo* passaging (Jungwirth et al., 2018) (**Supplementary Figure S1a**). D2A1 metastatic cells showed increased ACLY and FASN expression when compared to parental cells (**Supplementary Figure S1b**).

Although maintaining a high rate of *de novo* FA synthesis concomitantly to lipid breakdown (i.e., fatty acid beta-oxidation, FAO) could be biochemically and energetically unproductive, it has been reported that this apparently futile metabolic loop can occur in breast cancer cells that survive tumor regression (Havas et al., 2017). To investigate the nutrient contribution to the oxidative metabolism of TNBC cells, we real-time measured the Oxygen Consumption Rate (OCR) in 67NR, 4T07, and 4T1 cells using Seahorse analysis (**Supplementary Figures S1c,d**). Mitochondrial Stress Test was performed either in the presence of a mitochondrial pyruvate carrier inhibitor (UK5099), of a glutaminase inhibitor (BPTES), or by blocking lipid entry into the mitochondria using the CPT1a inhibitor etomoxir. This approach allows for dissecting the oxidative contribution of glucose-derived pyruvate, glutamine, and FA, respectively. Although UK5099 had the maximal OCR inhibition in the three cell lines (i.e., glucose dependency, **Supplementary Figure S1c**), etomoxir significantly and selectively impaired basal and maximal respiration in 4T07 and 4T1 (**Supplementary Figure S1d**), a dependence that was validated using the Oroboros oxygraph-2K high-resolution respirometer (**Supplementary Figure S1e**), indicating that the catabolic contribution of lipids is higher in the more aggressive models.

### Aberrant lipid metabolism sensitizes aggressive TNBC cells to ferroptosis

To identify potential metabolic vulnerabilities associated with the lipid metabolic reprogramming displayed by the more aggressive TNBC cells, we comparatively assessed the changes in cell viability induced in 67NR and 4T1 cells exposed to agents that target lipid-related metabolic pathways. Unexpectedly, targeting ACLY (SB204990), Acetyl-CoA Carboxylase (ACC, TOFA), FASN (TVB3166), FAO (etomoxir), LD biogenesis (Diacylglycerol Acyltransferase 1 and 2, DGAT1 and DGAT2, inhibitors) or lipid mobilization from LD (Adipose Triglyceride Lipase, ATGL, inhibitor, ATGListatin) showed no significant difference between 67NR and 4T1 cells (**Figure 2a**). Based on this evidence, we hypothesized that the lipid metabolism of the more aggressive cells does not impact the total amount of intracellular lipids but rather their composition and complexity, a proxy of desaturases activity. Indeed, the production of MUFA is catalyzed by SCD, whereas FADS are responsible for subsequent double bonds insertion and PUFA formation (Peck & Schulze, 2016). Western blot analysis revealed increased expression levels of SCD1, SCD2, FADS1, and FADS2 in the 4T07 and 4T1 cells (**Figure 2b**). A similar trend was observed at protein or mRNA level in the D2A1 series (**Supplementary Figure S2a**). However, the administration of either SC-26196, a selective FADS2 inhibitor, CP-24879, a FADS1/FADS2 dual inhibitor, or CAY10566, a SCD1 inhibitor, at concentrations able to inhibit the desaturases activity and not be *per se* being antioxidant (Lee et al., 2020), did not affect cell survival of 4T07 and 4T1 cells (**Supplementary Figure S2b**).

**Figure 2.**
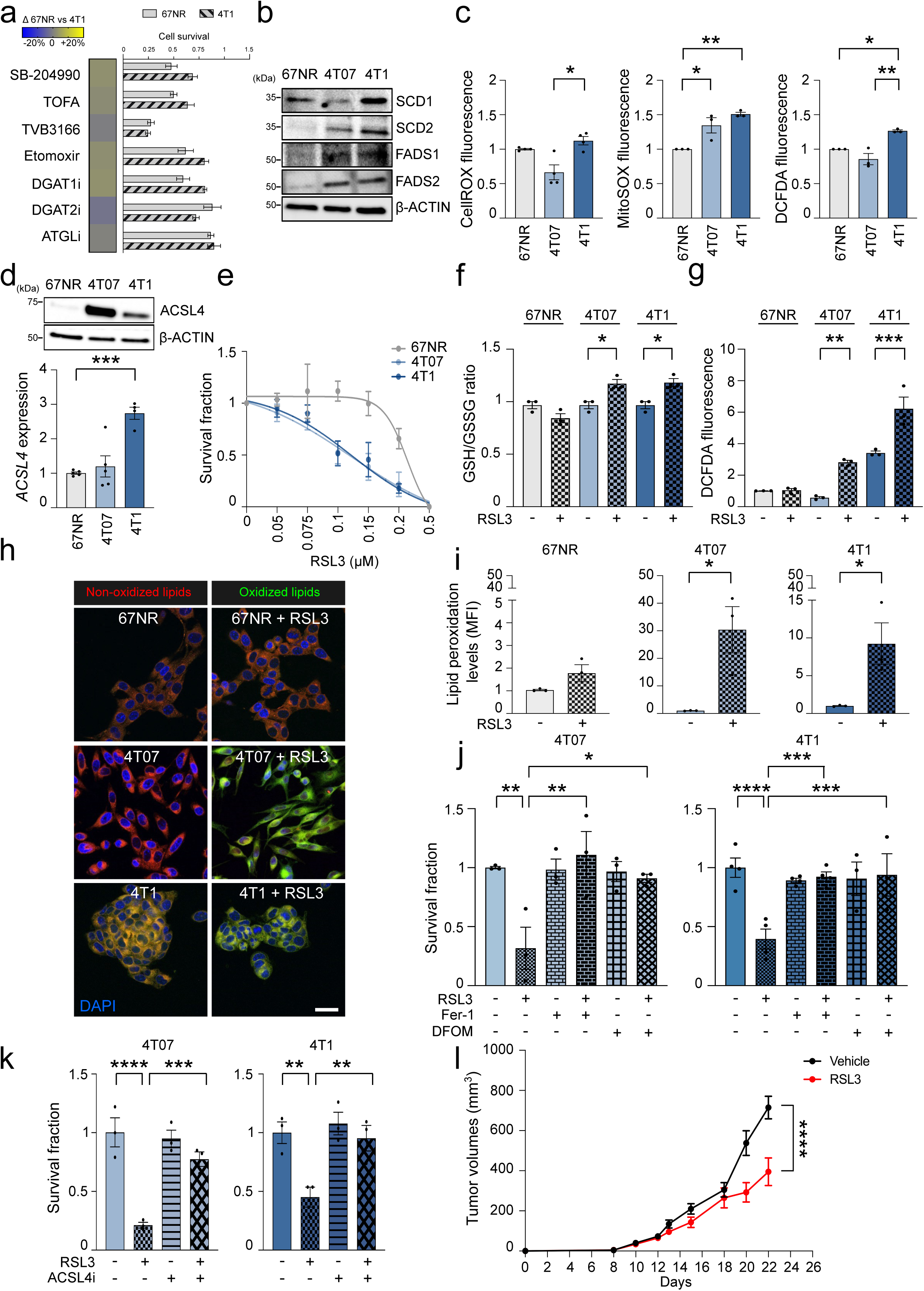
Aberrant lipid metabolism sensitizes aggressive TNBC cells to ferroptosis. (a) 67NR and 4T1 cells were treated with a series of drugs targeting lipogenic enzymes and subjected to cell viability assays. Heatmap shows that no drug exerts a differential effect between less aggressive and metastatic cells. Data represent means ± SEMs. Student’s t test. (b) Murine breast cancer cells were analyzed by WB analysis using the antibodies described in the figure. (c) Intracellular ROS levels were measured by CellROX and 2’,7’-dichlorofluorescin diacetate (DCFDA) staining while mitochondrial ROS levels by MitoSOX in TNBC cells. The non-metastatic 67NR cells were used as comparator. Data represent means ± SEMs. One-way ANOVA; Dunnett corrected. (d) TNBC cells were analyzed by qRT-PCR and WB analysis using the assay and the antibody described in the figure. Fold relative enrichment is shown using the non-metastatic cells as comparator. Data represent means ± SEMs. One-way ANOVA, Dunnett corrected. (e) 24-hour dose-response curve of RSL3 showed a differential effect between less aggressive and metastatic cells. Data represent means ± SEMs. One-way ANOVA, Bonferroni corrected. (f) GSH/GSSG concentration ratio was measured in TNBC cells treated with 0.1 μM RSL3 for 24 hours. Data represent means ± SEMs, n=3. One-way ANOVA, Bonferroni corrected. (g) Intracellular ROS levels were measured by DCFDA in TNBC cells treated with 0.1 μM RSL3 for 24 hours. The non-metastatic 67NR cells were used as comparator. Data represent means ± SEMs. One-way ANOVA; Dunnett corrected. (h,i) Lipid peroxidation level was evaluated by measuring the fluorescence intensity of BODIPY^581/591^ C11 using confocal microscopy (h) and cytofluorimetric analysis (i) in murine TNBC cells administrated with 1 μM of RSL3 for 2 hours. (Representative confocal images are shown. Oxidized lipids: green; non-oxidized lipids: red; nuclei: blue, DAPI; scale bar, 10 μm). Data represent means ± SEMs. One-way ANOVA; Dunnett corrected. (j) 4T07 and 4T1 cells were pre-treated with 15 μM Fer-1 or 5 μM DFOM for 4 hours and then administrated ON with 0.1 μM RSL3. After 24 hours cells were subjected to cell viability assay. One-way ANOVA; Dunnett corrected. (k) The indolent (4T07) and the highly (4T1) metastatic cells were pre-treated with 10 μM ACSL4i for 4 hours and then administrated ON with 0.1 μM RSL3. After 24 hours cells were subjected to cell viability assay. One-way ANOVA; Dunnett corrected. (l) *In vivo* response to RSL3 treatment in BALB/c mice. Mean±SEM (n=4-6 mice/group). Each dot represents a biological replicate. * *P*<0.05; ** *P*<0.01; *** *P*<0.001; **** *P*<0.0001.

Therefore, we postulated that impacting the availability of PUFA could be a metabolic vulnerability when aggressive cells are exposed to a secondary hit. Indeed, it is established that peroxidation of PUFA drives ferroptosis (Yang et al., 2016) and that MUFA have a ferroptosis-protective effect by suppressing lipid reactive oxygen species (ROS) accumulation at the plasma membrane (Magtanong et al., 2019). Notably, 4T07 and 4T1 cells showed higher cytosolic and/or mitochondrial ROS basal content compared to 67NR cells, as revealed by different redox-sensitive probes (**Figure 2c**), a trait that could be related to their enhanced oxidative metabolism (**Supplementary Figures S1c-e**). ACSL4 is crucial for PUFA-mediated ferroptosis execution, and its protein and mRNA expression levels are higher in the more aggressive cells of the 4T1 and D2A1 series (**Figure 2d and Supplementary Figure S2c**). This molecular, metabolic, and redox cellular asset could render the more aggressive TNBC cells highly susceptible to ferroptosis induction. We, therefore, subjected the 4T1 series (**Figure 2e**) and the D2A1 series (**Supplementary Figure S2d**) to increasing doses of the ferroptosis inducer RSL3 (RAS-selective lethal), which inhibits GPX4 (Yang et al., 2014). RSL3 reduced cell survival in the more aggressive cells in a dose-dependent manner, whereas the less aggressive 67NR and D2A1 were much more resistant to ferroptosis induction (**Figure 2e and Supplementary Figure S2d**). Since ROS production and subsequent lipid peroxidation are both hallmarks of ferroptosis, we evaluated upon RSL3 treatment (i) the ratio of reduced and oxidized glutathione levels (GSH/GSSG, **Figure 2f**), (ii) ROS content using DCFDA probe (**Figure 2g**) and (iii) lipid peroxidation using the fluorescent lipid peroxidation-sensitive dye BODIPY^581/591^-C11 by confocal and flow-cytometry analyses (**Figure 2h,i**). Altered GSH/GSSG proportion and increased ROS content were observed exclusively in the 4T07 and 4T1 cells. Although BODIPY^581/591^-C11 should be immediately oxidized after RSL3 administration, this did not occur in 67NR exposed to RSL3. Conversely, a strong increase in the oxidized form of BODIPY^581/591^-C11 was observed in the more aggressive cells exposed to RSL3 (**Figure 2h,i**). Timing (2 hours) and dosing (1 μM) were chosen not to induce cell death in cell lines. Similarly, RSL3 rapidly elicited a strong increase in the oxidized form of BODIPY^581/591^-C11 in the D2A1-m1 and D2A1-m2 cells (**Supplementary Figure S2e**). To exclude a restricted RSL3-mediated effect, the 4T1 and D2A1 series were exposed to erastin, a ferroptosis inducer that inhibits the cystine/glutamate antiporter (system XC^−^) known to provide cystine (i.e., the oxidized form of cysteine) required for glutathione synthesis (Dixon et al., 2012). Importantly, the more aggressive TNBC cells were sensitive to erastin administration (**Supplementary Figure S2f,g)** and displayed enhanced levels of lipid peroxides (**Supplementary Figure S2h-j**).

To finally confirm that aggressive TNBC cells undergo ferroptosis, we exposed the ferroptosis-sensitive 4T1, 4T07, D2A1-m1 and D2A1-m2 cells to RSL3 or erastin together with either the radical-trapping antioxidant ferrostatin-1 (Fer-1), the iron chelator agent deferoxamine (DFOM, **Figure 2j**, **Supplementary Figure S2k-m**) or an ACSL4 inhibitor (**Figure 2k**). All these ferroptosis-preventing approaches were sufficient to abrogate the cell death induced by both RSL3 and erastin.

To assess whether this metabolic vulnerability could be exploited *in vivo*, we used the syngeneic 4T1 model. 4T1 cells were subcutaneously injected in BALB/c mice where they formed palpable tumors after 7 days. Mice were then randomized and treated with RSL3 for 2 weeks. RSL3 significantly reduced tumor growth (**Figure 2l**) without showing severe toxicity (**Supplementary Figure S2n**).

### Targeting LD biogenesis potentiates the effect of RSL3 in TNBC cells

LD have a crucial role in tumor progression and are involved in ferroptosis protection (Danielli et al., 2023). Particularly, storing PUFA in LD reduces the availability of double bonds for peroxidation (Dierge et al., 2021). Since 4T1 cells showed higher LD content (**Figure 1f**) and higher FADS1/2 expression (**Figure 2b**), we monitored LD content upon FADS inhibition. Interestingly, SC-26196 and CP-24879 administration reduced LD content, assessed using BODIPY^493/503^ (**Figure 3a**), suggesting that the increased LD content of 4T1 cells was partially due to their ability to synthesize PUFA. Crucially, RSL3 administration resulted in a significant and rapid increase in the LD content of the more aggressive models (**Figure 3b,c**), suggesting a potential scavenging mechanism exploited by the cells in an attempt to resist the ferroptosis-promoting insult. Conversely, no change was observed in the RSL3 insensitive 67NR. Although 4T07 and 4T1 cells were already sensitive to RSL3, impairing LD biogenesis using the DGAT1 and DGAT2 inhibitors potentiated the cell death induced by RSL3, particularly in the more aggressive 4T1 cells (**Figure 3d**). This synthetic lethal approach could be further investigated in those ferroptosis-resistant cells that are characterized by increased LD content.

**Figure 3.**
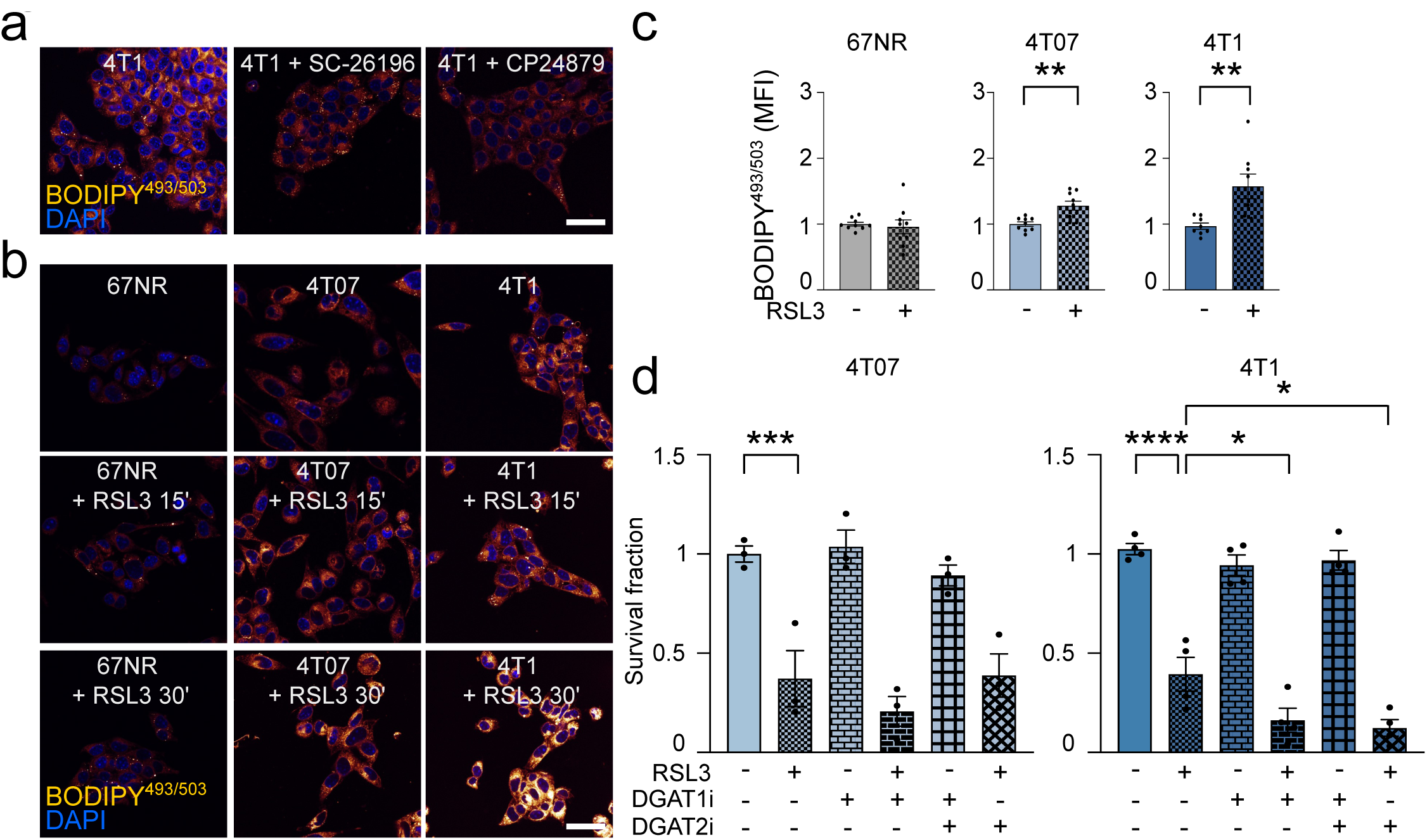
Targeting LD biogenesis potentiates the effect of RSL3 in TNBC cells. (a) 4T1 cells treated for 24 hours with 10 μM FADS2i or 10 μM FADS1/2i were subjected to BODIPY^493/503^ staining by confocal microscopy. Representative images are shown (orange yellow: LD; blue: DAPI, nuclei; scale bar, 10 μm). Data represent means ± SEMs. LD numbers were quantified as described in Methods. (b,c) TNBC cells treated with 1 μM RSL3 for 15’ and 30’ were subjected to confocal (b) and cytofluorimetric (c) analyses. Representative pictures of BODIPY^493/503^ stained cells are shown (orange/yellow: LD; blue: DAPI, nuclei). Scale bar, 10 μm. Data represent means ± SEMs. One-way ANOVA; Dunnett corrected. (d) The indolent (4T07) and the highly (4T1) metastatic cells were pre-treated with 10 μM DGAT1i or DGAT2i for 4 hours and then administrated ON with 0.1 μM RSL3. After 24 hours cells were subjected to cell viability assay. One-way ANOVA; Dunnett corrected. Each dot represents a biological replicate. * *P*<0.05; ** *P*<0.01; *** *P*<0.001; **** *P*<0.0001.

### Ferroptosis induction alters the lipidomic profile of sensitive TNBC cells but not that of the resistant cells

To explore the potential lipidic alterations induced by ferroptosis execution in sensitive and resistant cells, we subjected 67NR (resistant) and 4T1 (sensitive) cells, either untreated or exposed to a short-acute RSL3 treatment, to lipid extraction and subsequent lipidomic analysis (Cattaneo et al., 2021). Partial Least Square Discriminant Analysis (PLS-DA) of 1209 unambiguously identified lipids showed a clear separation between 4T1 cells and 4T1 exposed to RSL3 (**Figure 4a** and **Supplementary Table 1**). Conversely, 67NR exposed or not to RSL3 did not cluster apart in the PLS-DA, suggesting very few potential differences in their lipidomic profile (**Figure 4a**). Indeed, the analysis of lipid entities differentially expressed (adj. t-test; P<0.05, fold-change=1.5) upon RSL3 exposure identified 47 over-represented and 66 under-represented lipids in the 4T1 cells treated with RSL3 compared to untreated cells (**Figure 4b**), whereas only 1 and 12 entities were over and under-represented in the 67NR pair (**Figure 4c**), suggesting that RSL3 was neither able to induce cell death in the 67NR nor impact on their lipidomic profile. It is reported that FADS expression and activity determine ferroptosis sensitivity in gastric and ovarian cancer (Lee et al., 2020; Xuan et al., 2022), whereas those of SCD sustain ferroptosis resistance in breast cancer (Luis et al., 2021), reinforcing the idea that PUFA intracellular availability is necessary for ferroptosis execution. In mammalian membranes, PUFA within the phospholipids are mainly phosphatidylcholine (PC) and phosphatidylethanolamine (PE) (van der Veen et al., 2017). By analyzing the lipidomic profiles, the PUFA/MUFA ratio of PC and their derivatives lysophosphatidylcholine (LPC) were significantly higher in the 4T1 cells than in 67NR, and a similar trend was also observed for lysophosphatidylethanolamine (LPE) (**Figure 4d**), a trait that explains the susceptibility of 4T1 cells to ferroptosis-inducing insults.

**Figure 4.**
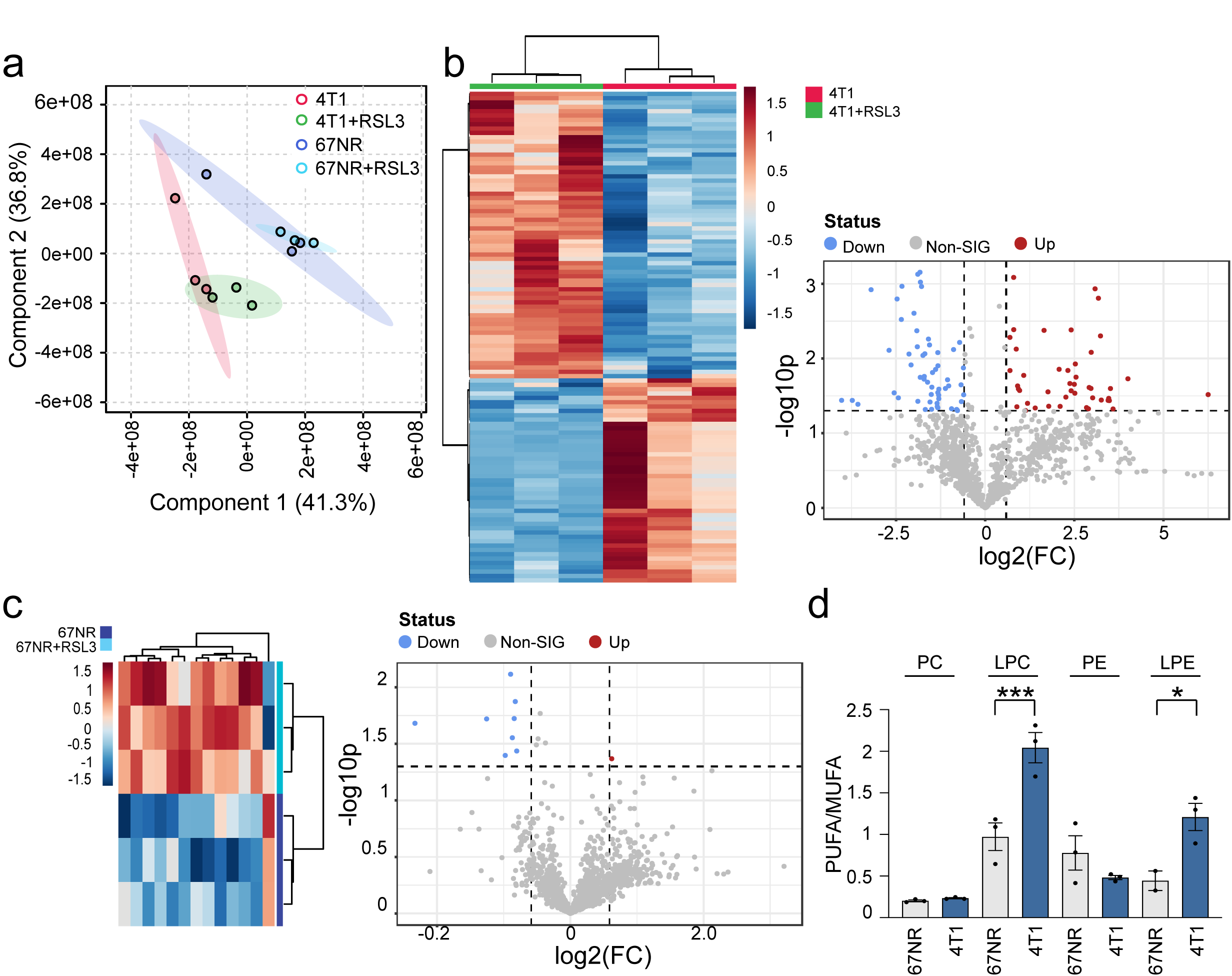
Ferroptosis induction alters the lipidomic profile of sensitive TNBC cells but not that of the resistant cells. (a) Partial Least Square Discriminant analysis (PLS-DA) of the lipidomic profile of 67NR and 4T1 cells with and without 1 μM RSL3 for 2 hours. (b) Heatmap of the significantly and differentially expressed lipid entities of the 4T1 cells exposed to 1 μM RSL3 for 2 hours (top). The data were represented as volcano plot (bottom), highlighting the upregulated (red) and downregulated (blue) lipid species (fold change threshold of 1.5 and p-value <0.05). (c) Heatmap of the significantly and differentially expressed lipid entities of the 67NR cells exposed to 1 μM RSL3 for 2 hours (top). The data were represented as volcano plot (bottom), highlighting the upregulated (red) and downregulated (blue) lipid species (fold change threshold of 1.5 and p-value <0.05). (d) Ratio of PUFA/MUFA in 4T1 and 67NR cells for PC, LPC, PE and LPE lipid classes, calculated as described in the Material and Methods section. Student t-test. Each dot represents a biological replicate. * *P*<0.05; *** *P*<0.001.

### Targeting FADS1/2 prevents ferroptosis induction in aggressive TNBC

To demonstrate whether FADS1 and FADS2 have a role in the ferroptotic cell death, we first exposed the ferroptosis-sensitive cells to SC-26196 (FADS2 inhibitor) and CP-24879 (FADS1/FADS2 dual inhibitor). The inhibition of FADS1 and 2 prevented the RSL3- (**Figure 5a and Supplementary Figure S3a**) and the erastin- (**Supplementary Figure S3b,c**) induced cell death in 4T07, 4T1, D2A1-m1 and D2A1-m2 cells. Conversely, the inhibition of SCD1 was not sufficient to prevent ferroptosis (**Figure 5a**). Confocal and flow-cytometry analyses of the oxidized form of BODIPY^581/591^-C11 confirmed the prevention of lipid peroxidation exerted by FADS inhibition in 4T1 and 4T07 cells (**Figure 5b,c and Supplementary Figure S3d,e**) and in D2A1-m1 and D2A1-m2 cells (**Supplementary Figure S3f**). Since no selective FADS1 inhibitors are available and CP-24879 is a FADS1/FADS2 dual inhibitor, we silenced FADS1 in 4T1 cells. FADS1 silencing (**Figure 5d**) prevented RSL3-induced ferroptosis (**Figure 5e**) and lipid peroxidation (**Figure 5f,g**) in 4T1 cells. To validate these observations, these assays were extended to two human TNBC cell lines that have different metastatic abilities, being the BT20 cells able to colonize distant sites but not to expand and the MDA-MB-231 highly metastatic and penetrant (Jin et al., 2020). Interestingly, MDA-MB-231 cells showed increased protein levels of ACSL4 and higher sensitivity to RSL3- and erastin-induced cell death (**Supplementary Figure S3g,i**), that Fer-1 administration was able to prevent (**Supplementary Figure S3h**), hence confirming that MDA-MB-231 cells are ferroptosis-sensitive. Moreover, MDA-MB-231 cells show higher expression of FADS1 and FADS2 compared to BT20 (**Supplementary Figure S3i**). As expected, both SC-26196 and CP-24879 prevented the RSL3- and erastin-induced cell death in MDA-MB-231 cells (**Supplementary Figure S3j,k**).

**Figure 5.**
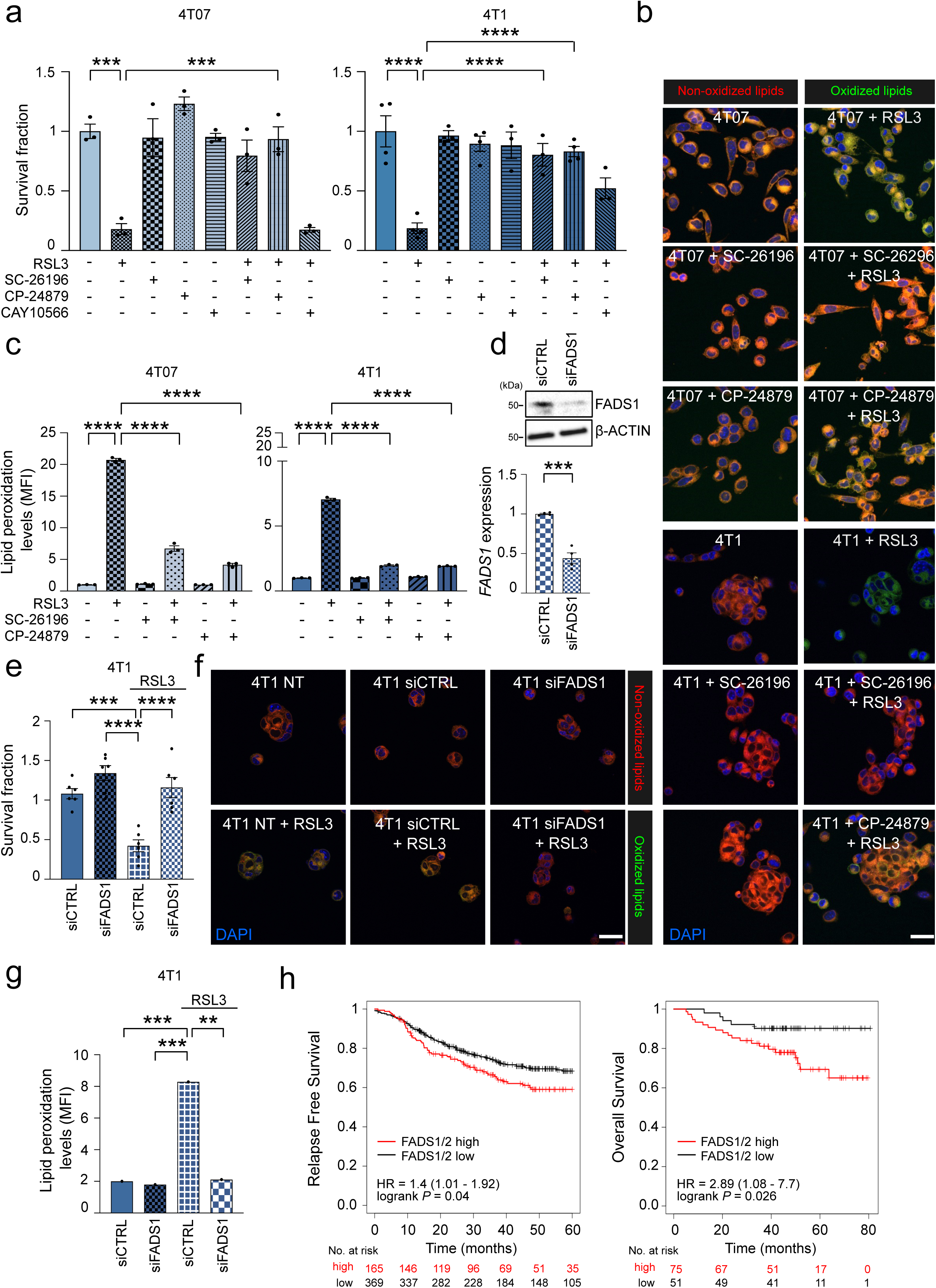
Targeting FADS1/2 prevents ferroptosis induction in aggressive TNBC. (a) TNBC metastatic cells were pre-treated with 10 μM FADS2i or FADS1/2i for 4 hours, administrated ON with 0.1 μM RSL3 and subjected to cell viability assay. One-way ANOVA; Dunnett corrected. (b,c) TNBC indolent and highly metastatic cells were pre-treated with 10 μM FADS2i or FADS1/2i for 4 hours, administrated ON with 0.1 μM RSL3 and subjected to confocal analysis (b) and cytofluorimetric analysis (c) to measure lipid peroxidation. Representative pictures of BODIPY^581/591^ C11 stained cells are shown (oxidized lipids: green; non-oxidized lipids: red; nuclei: blue, DAPI). Scale bar, 10 μm. Data represent means ± SEMs. One-way ANOVA; Dunnett corrected. (d-f) 4T1 cells were transfected with non-targeting small interfering RNA (siRNA) (siCTR) or siRNA targeting FADS1 (siFADS1) and subjected to qRT-PCR and WB analysis for FADS1 expression (d), administrated ON with 0.1 μM RSL3 and subjected to cell viability assay using crystal violet (e) or confocal analysis (f) and cytofluorimetric analysis (g) to measure lipid peroxidation. Representative pictures of BODIPY^581/591^ C11 stained cells are shown (oxidized lipids: green; non-oxidized lipids: red; nuclei: blue, DAPI). siCTR transfected cells were used as comparator. Data represent means ± SEMs. Student’s t test; One-way ANOVA; Dunnett corrected. (h) Kaplan-Meier analysis of relapse free survival (RFS) and overall survival (OS) of a curated cohort of TNBC patients divided into high and low as described in Material and Methods section for FADS1/FADS2 expression. HR and log-rank Mantel-Cox p values are shown. Each dot represents a biological replicate. *** *P*<0.001; **** *P*<0.0001.

The clinical relevance of these findings was validated retrospectively in a cohort of TNBC. TNBC-bearing patients were dichotomized based on higher and lower mRNA expression of *FADS1*, *FADS2,* or both (i.e., FADS1^high^ and FADS2^high^ expressing patients). High expression of FADS1 (hazard ratio [HR] = 1.39, log-rank P = 0.044, n = 534, and HR = 1.96, log-rank P = 0.1, n = 126, **Supplementary Figure S3l**), high expression of FADS2 (HR = 1.39, log-rank P = 0.064, n = 534, and HR = 2.31, log-rank P = 0.12, n = 126, **Supplementary Figure S3m**), and high expression of both desaturases (HR = 1.4, log-rank P = 0.04, n = 534, and HR = 2.89, log-rank P = 0.026, n = 126, **Figure 5h**) correlate with reduced survival and increased relapse in TNBC patients. These findings demonstrate that FADS1/2 expression has a prognostic value and identifies a subset of TNBC more susceptible to undergo ferroptosis.

### Exogenous FA supplementation alters ferroptosis execution in TNBC cells

In line with the major role of PUFA availability in sustaining ferroptosis execution in TNBC cells, we exposed the ferroptosis-resistant 67NR cells to an exogenous FA supplementation. Adrenic acid (C22:4; all-cis-7,10,13,16-docosatetraenoic acid), erucic acid (C22:1; cis-13-docosenoic acid), and palmitoleic acid (C16:1; cis-9-hexadecenoic acid) were individually added to 67NR cell medium either alone or concomitantly to RSL3. In line with the hypothesis, only the administration of the PUFA adrenic acid sensitized 67NR to RSL3 exposure, as revealed by increased cell death (**Figure 6a**) and lipid peroxidation (**Figure 6b**), whereas MUFA (C22:1 and C18:1) administration did not sensitize 67NR to RSL3. Importantly, the supplementation of 22:4, but not that of 22:1, enhanced RSL3-induced cell death in the more aggressive cells (**Figure 6c**), reinforcing the concept that PUFA availability is essential for ferroptosis execution while MUFA exert a protective effect in TNBC cells.

**Figure 6.**
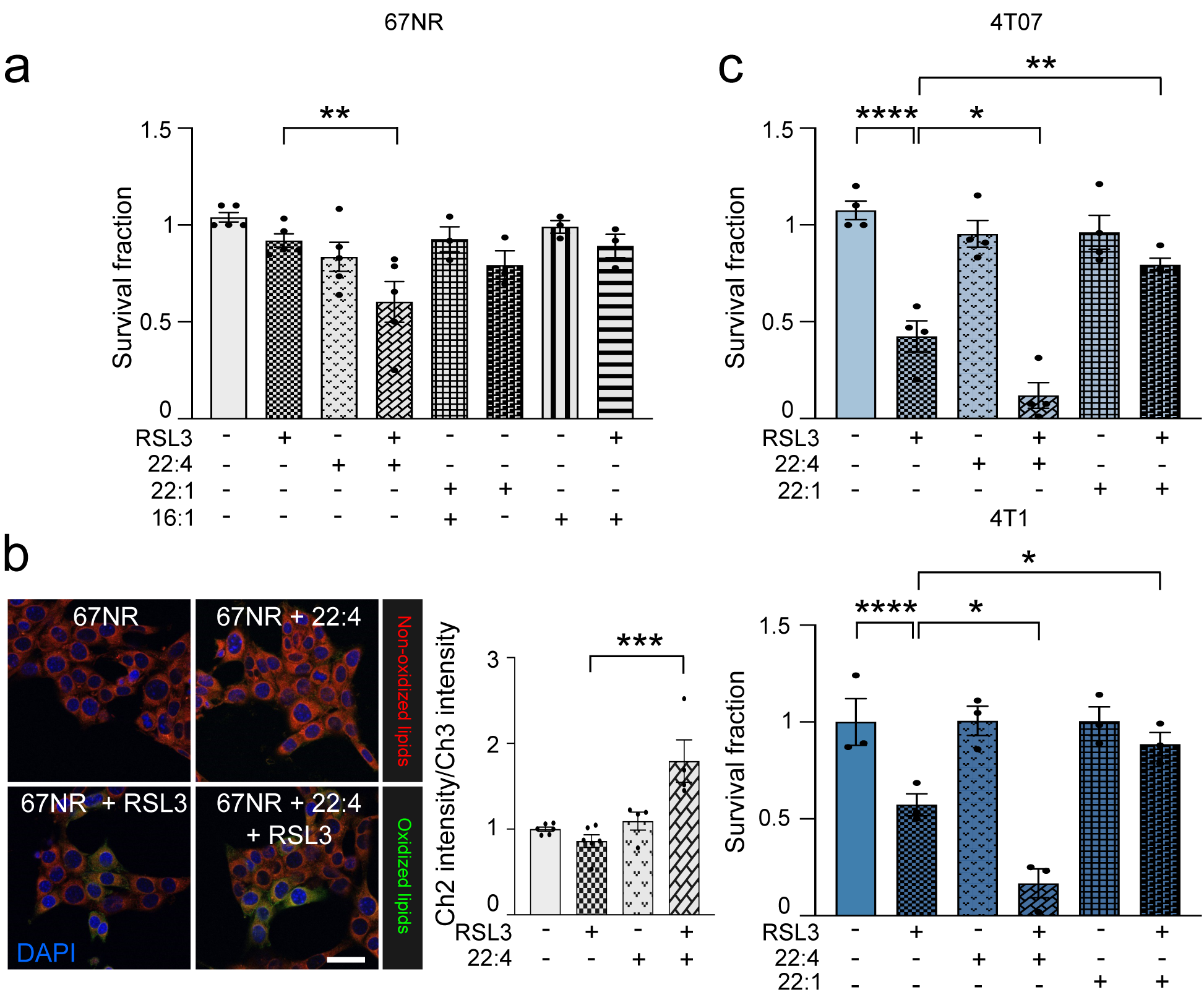
Exogenous FA supplementation alters ferroptosis execution in TNBC. (a) The non-metastatic 67NR cells were pre-treated with 10 μM 22:4, 10 μM 22:1, or 5 μM 16:1 for 4 hours and then administrated ON with 0.1 μM RSL3. After 24 hours cells were subjected to cell viability assay. One-way ANOVA; Dunnett corrected. (b) TNBC 67NR cells were pre-treated with 10 μM 22:4 for 4 hours and then administrated for 2 hours with 1 μM RSL3 and subjected to confocal analysis to measure lipid peroxidation. Representative pictures of BODIPY^581/591^ C11 stained cells are shown (oxidized lipids: green; non-oxidized lipids: red; nuclei: blue, DAPI). Quantification of BODIPY581/591 C11 spots was reported. Scale bar, 10 μm. Data represent means ± SEMs. One-way ANOVA; Dunnett corrected. (c) TNBC metastatic cells were pre-treated with 10 μM 22:4 or 22:1 for 4 hours, administrated ON with 0.1 μM RSL3 and subjected to cell viability assay. One-way ANOVA; Dunnett corrected. Each dot represents a biological replicate. * *P*<0.05; ** *P*<0.01; *** *P*<0.001; **** *P*<0.0001.

## Discussion

This study identifies a novel lipid metabolic reprogramming as a potential metabolic vulnerability that can be exploited in a subset of aggressive TNBC. Indeed, we have shown that TNBC characterized by increased FADS1/2 expression are susceptible to ferroptosis-inducing agents. This is crucial in addressing a biological and clinical need as it may allow stratifying patients that could benefit from a tailored approach in a subset of breast cancers in which therapy remains a challenge.

A recent integrated multi-omics analysis of a large cohort of TNBC reported heterogeneous phenotypes in ferroptosis-related metabolites and metabolic pathways (Yang et al., 2023). Particularly, the luminal androgen receptor (LAR) TNBC subtype was characterized by ROS accumulation, enriched FA metabolism and lipogenic phenotype, increased levels of oxidized phosphatidylethanolamines, and deregulation of glutathione metabolism, which render LAR TNBC sensitive to GPX4 inhibitors (Yang et al., 2023). Although the authors reported increased levels of intracellular PUFA in ferroptosis-sensitive LAR TNBC, they did not prove that this was due to increased PUFA biosynthesis (i.e., FADS1/2 activity). However, in other cancer types, the increased expression levels and subsequent activity of the enzymatic asset involved in the formation of long and unsaturated FA is a major determinant of ferroptosis sensitivity. Lee *et al*. reported that mesenchymal-type gastric cancers, but not those of the intestinal subtype, are sensitive to the GPX4 inhibitors RSL-3 and ML-210 (Lee et al., 2020). An in-depth metabolic and metabolomic profiling of this ferroptosis-sensitive subset revealed increased intracellular levels of PUFA, particularly phosphatidylethanolamine (PE)-linked arachidonic acid and adrenic acid, which are synthesized in cancers with high expression levels of the ELOVL5 (elongation of very long chain fatty acids protein 5) and FADS1 (Lee et al., 2020). Moreover, Xuan *et al*. reported increased levels and activity of SCD1 and FADS2 in metastatic ovarian cancers (Xuan et al., 2022).

However, the intracellular levels of oxidable PUFA in ferroptosis-sensitive cells are also dependent on extracellular availability and exposure to exogenous FA, since it is established that altering the intracellular relative composition between MUFA and PUFA by exogenous FA supplementation is essential for ferroptosis execution (Magtanong et al., 2019). Indeed, Ubellacker *et al*. showed an anti-ferroptosis effect of oleic acid supplementation in melanoma (Ubellacker et al., 2020) and Dierge *et al*. showed that cancer cells can undergo ferroptosis in an acidic environment when supplemented with PUFA but not with oleic acid (Dierge et al., 2021). Mechanistically, the authors showed that preventing LD formation upon PUFA supplementation did not protect cancer cells from the enhanced uptake of PUFA and subsequent lipid peroxidation, particularly of those PUFA incorporated into the plasma membrane. In line with these reports, we herein demonstrated that PUFA administration sensitized the less aggressive TNBC cells to ferroptosis and potentiated the effects of the ferroptosis-inducing agents in the aggressive models, whereas MUFA supplementation has the opposite effects.

The importance of the crosstalk between LD and ferroptosis susceptibility emerged also in our study. Indeed, when aggressive TNBC cells are exposed to ferroptosis-inducing agents, they immediately activate a “defensive” mechanism involving a rapid LD formation. However, our data showed that aggressive TNBC cells are characterized by high basal levels of LD, a feature that is paralleled by increased ROS levels and lipid biosynthesis. It is reasonable to postulate that, although LD turnover can be activated in these cells, it is not sufficient to buffer the deleterious effects induced by RSL3 that concomitantly influence lipids availability and redox homeostasis. An important aspect that was not addressed by our study is why increasing desaturases expression might be advantageous for aggressive TNBC. There could be several explanations since (i) enhanced PUFA within the plasma membrane could facilitate cell movement and therefore be linked to the increased invasive potential of these cancers (Harayama & Antonny, 2023), as it could be in our models; (ii) eicosanoids, a class of bioactive signaling lipids, are synthesized from arachidonic acid and have important pro-tumoral signaling role (Koundouros et al., 2020; Wang & Dubois, 2010); (iii) PUFA (independently of the source, i.e., also released by cancer cells) can promote a pro-inflammatory tumor microenvironment, which is known to promote tumor initiation and progression (Marion-Letellier et al., 2015; Vriens et al., 2019).

In conclusion, our data show that lipid metabolism can reveal metabolic vulnerabilities within a subset of breast cancers that lack prognostic markers and selective therapeutic strategies. Importantly, many preclinical studies have proposed to target lipid metabolism, but the clinical translation has been so far challenging and ineffective. Ferroptosis-inducing agents have been proposed in different tumor contexts although so far this approach has been tested in preclinical models. However, dietary approaches that influence iron availability (Delesderrier et al., 2023) and selenocysteine synthesis (Chen et al., 2013), two major players involved in ferroptosis execution, could have a more rapid therapeutic translation and therefore, identifying those TNBC patients that are ferroptosis sensitive (or resistant) may be of clinical relevance. Finally, since FA supplementation alters cancer ferroptosis sensitivity, dietary approaches will be essential for potentiating the efficacy of a pro-ferroptosis approach not only in ferroptosis-sensitive tumors but also in those that are intrinsically resistant. In addition, since ferroptosis execution is immunogenic (Yang et al., 2023), ferroptosis-promoting agents can be investigated prospectively not only as monotherapy but in combination with immunotherapy or other anti-tumoral drugs (Minami et al., 2023). Impacting on the cancer desaturases activity (whether expressed by the cancer cells or by other tumor-associated stromal cells) could therefore change the FA composition of the tumor microenvironment and influence the success of the ferroptosis-inducing approach.

## Acknowledgment

The work was funded by *Associazione Italiana Ricerca sul Cancro* (AIRC) and *Fondazione Cassa di Risparmio di Firenze* (grant Multiuser 19515 to PC and AM) and AIRC (grant IG 22941 to AM, grant IG 8797 to PC). NL is supported by an AIRC fellowship. AS is supported by *Associazione Annastaccatolisa*. LI is supported by *Fondazione Pezcoller*. We thank Prof Clare Isacke (The Institute of Cancer Research, London) for the fruitful discussion and insightful comments. The data presented in the current study were in part generated using the equipment of the *Facility di Medicina Molecolare*, funded by “*Ministero dell’Istruzione dell’Università e della Ricerca – Bando Dipartimenti di Eccellenza 2018-2022*”. The illustration was created with BioRender.com.

## Author contributions

The authors contributed to this work in different capacities, described as follows. NL, AS performed cell-based and biochemistry experiments, study design, data acquisition, analysis, and interpretation and wrote the manuscript; AS, MB, FB performed cell-based experiments, *in silico* analysis and participated in the interpretation of the data; AC, DLL, MI, AA performed *in vivo* based experiments and subsequent specimens derived analysis; LI, GC, EG, PC, IM performed the data analysis and interpretation and revised the manuscript, LT, AB performed the lipidomic experiments and subsequent computational analysis; AM conceived, designed, and supervised the study; analyzed and interpreted the data; and wrote the original draft of the manuscript. All the authors reviewed the prepared manuscript.

---

**Supplementary Figure 1.**
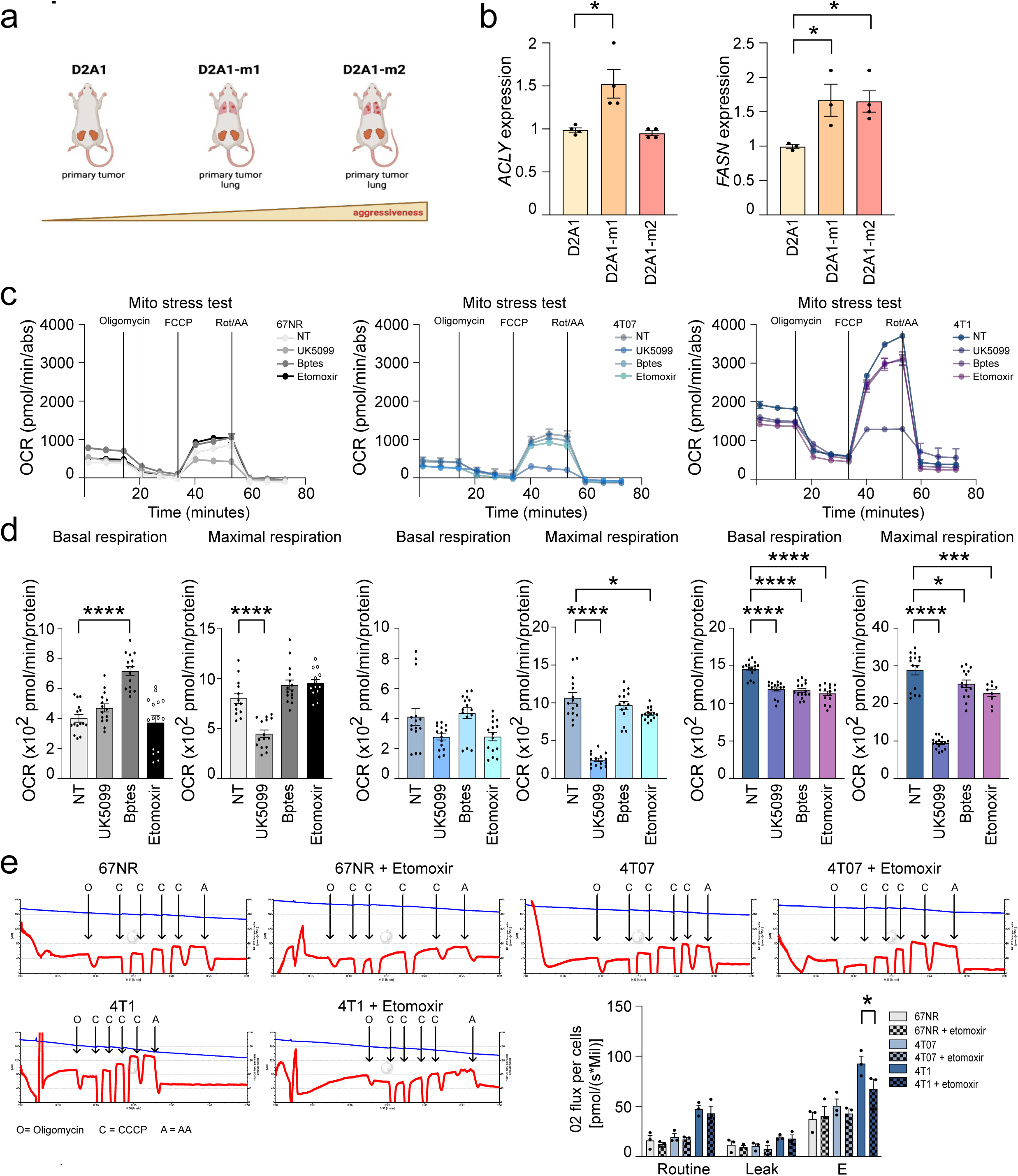
(a) Schematic representation of the metastatic potential of the D2A1 cell line series. (b) Murine breast cancer D2A1 cells and their metastatic derivatives D2A1-m1 and D2A1-m2 were analyzed by qRT-PCR analysis using the assays described in the figures. Fold relative enrichment is shown using the weakly metastatic parental cells as comparator. Data represent means ± SEMs. One-way ANOVA, Dunnett corrected. (c-d) Seahorse XFe96 Mito Stress Test was performed on 4T1 cell line series treated with 2 μM UK5099, 3 μM BPTES, or 4 μM Etomoxir for 30 minutes in the presence of standard condition (full medium), and oxygen consumption rate (OCR) was calculated in real-time after the administration of the ATP synthase inhibitor oligomycin (oligo), the proton uncoupler carbonilcyanide p-triflouromethoxyphenylhydrazone (FCCP), and the respiratory complex I inhibitor rotenone together with the respiratory complex III inhibitor antimycin A (Rot/AA) (c). Basal and maximal respiration was calculated as described in Methods and normalized on protein content (d). Each dot represents at least five technical replicates from three biological replicates. Data represent means ± SEMs. One-way ANOVA; Dunnett corrected. (e) TNBC cells were treated 40 μM of Etomoxir for 30 minutes. After detachment, cells were subjected to high-resolution respirometry analysis by Oroboros-O2K instrument. Representative graphs of cell respirometry analysis in the control (left) and treatment (right) conditions. The blue curve represents the oxygen concentration, whereas the red slope shows the oxygen consumption before and after the serial injections of oligomycin (Oligo), uncoupler CCCP, and Antimycin (AA). (e) Bar chart graph of basal oxygen consumption (ROUTINE), proton leak (LEAK) and maximal oxygen consumption (E) values subtracted from residual oxygen consumption (ROX) in control and Etomoxir treated cells. Data represent means ± SEMs. One-way ANOVA; Dunnett corrected. Each dot represents a biological replicate. * *P*<0.05; ** *P*<0.01; *** *P*<0.001; **** *P*<0.0001.

**Supplementary Figure 2.**
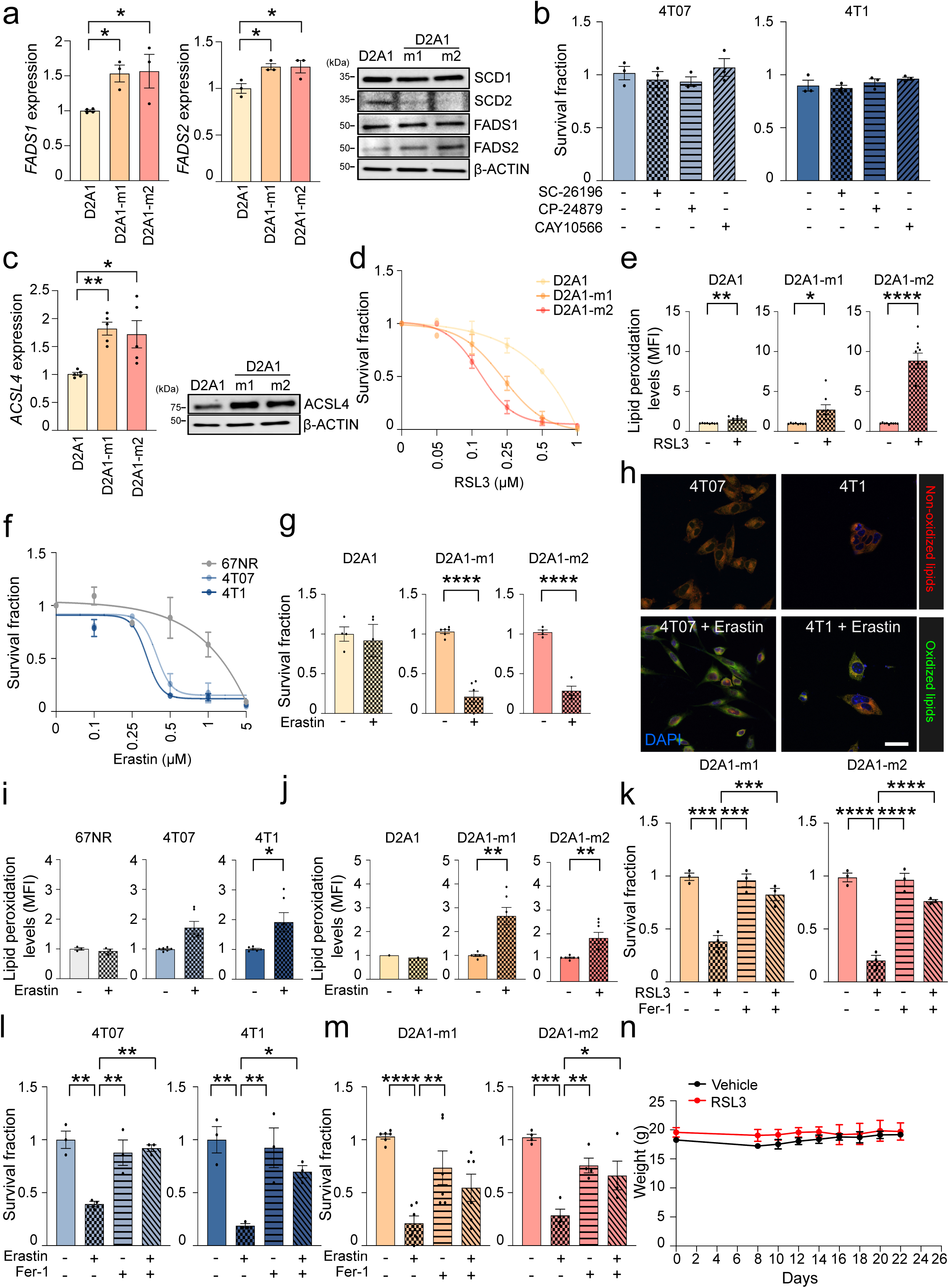
(a) D2A1 cell line series were analyzed by qRT-PCR and WB analysis using the assays and the antibodies described in the figure. Fold relative enrichment is shown using the less metastatic cells as comparator. Data represent means ± SEMs. One-way ANOVA, Dunnett corrected. (b) TNBC metastatic cells were treated with 10 μM FADS2i, FADS1/2i or SCD1i for 4 hours, administrated ON with 0.1 μM RSL3 and subjected to cell viability assay. One-way ANOVA; Dunnett corrected. (c) Murine breast cancer cells were analyzed by qRT-PCR and WB analyses using the assay and the antibody described in the figures. Fold relative enrichment is shown using the weakly metastatic parental cells as comparator. Data represent means ± SEMs. One-way ANOVA, Dunnett corrected. (d) 24-hour dose-response curve of RSL3 showed a differential effect between less aggressive and metastatic D2A1 series cells. Data represent means ± SEMs. One-way ANOVA, Bonferroni corrected. (e) Lipid peroxidation level was evaluated by measuring the fluorescence intensity of BODIPY^581/591^-C11 using cytofluorimetric analysis in murine TNBC cells administrated with 1 μM of RSL3 for 2 hours. (f) 24-hour dose-response curve of erastin showed a differential effect between weakly and highly aggressive 4T1 series cells. Data represent means ± SEMs. One-way ANOVA, Bonferroni corrected. (g) D2A1 cell lines series were treated with 0.5 μM erastin for 24 hours and subjected to cell viability assay. (h-i) Lipid peroxidation level was evaluated by measuring the fluorescence intensity of BODIPY^581/591^-C11 using confocal microscopy (h) and cytofluorimetric analyses (i) in murine TNBC cells administrated with 5 μM of erastin for 2 hours. (Representative confocal images are shown. Oxidized lipids: green; non-oxidized lipids: red; nuclei: blue, DAPI; scale bar, 10 μm). Data represent means ± SEMs. One-way ANOVA; Dunnett corrected. (j) The D2A1 cell line series administrated with 5 μM of erastin for 2 hours was subjected to cytofluorimetric analysis. FACS analysis of the MFI of the populations positive for BODIPY^581/591^-C11 was reported. Data represent means ± SEMs. One-way ANOVA, Dunnett corrected. (k) D2A1-m1 and D2A1-m2 cells were pre-treated with 15 μM Fer-1 or 5 μM DFOM for 4 hours and then administrated ON with 0.1 μM RSL3. After 24 hours cells were subjected to cell viability assay. One-way ANOVA; Dunnett corrected. (l) TNBC metastatic cells were pre-treated with 15 μM Fer-1 or 5 μM DFOM for 4 hours and then administrated ON with 0.5 μM erastin. After 24 hours cells were subjected to cell viability assay. One-way ANOVA; Dunnett corrected. (m) D2A1 metastatic cells were pre-treated with 15 μM Fer-1 or 5 μM DFOM for 4 hours and then administrated ON with 0.5 μM erastin. After 24 hours cells were subjected to cell viability assay. One-way ANOVA; Dunnett corrected. (n) Weight of BALB/c mice exposed for 15 days to 40mg/kg RSL3. Data represent means ± SEMs. Student’s t test; One-way ANOVA; Each dot represents a biological replicate. * *P*<0.05; ** *P*<0.01; *** *P*<0.001; **** *P*<0.0001.

**Supplementary Figure 3.**
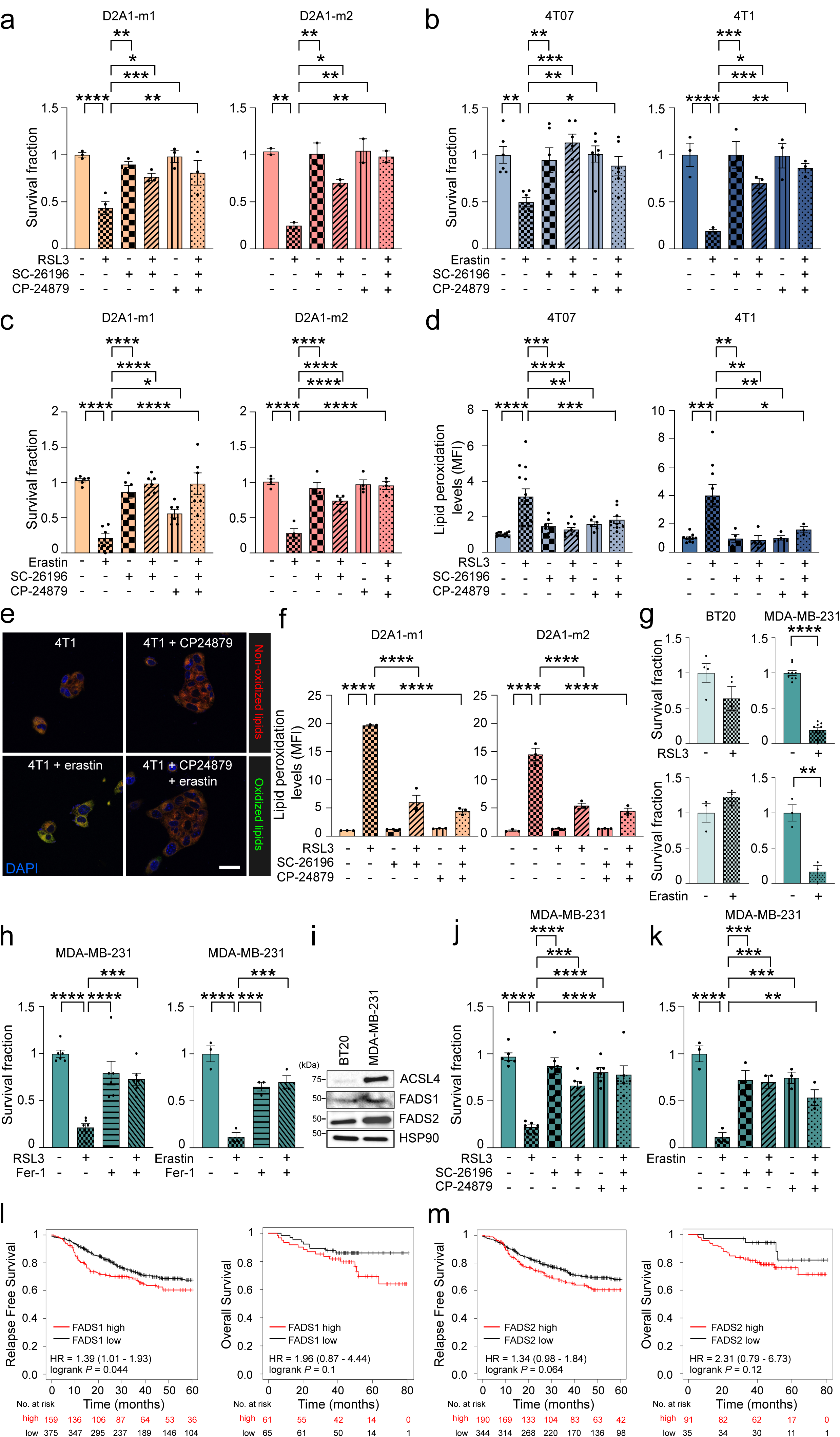
(a) D2A1-m1 and D2A1-m2 metastatic cells were pre-treated with 10 μM FADS2i or FADS1/2i for 4 hours, administrated ON with 0.1 μM RSL3, and subjected to cell viability assay. One-way ANOVA; Dunnett corrected. (b-c) 4T07, 4T1 (b), D2A1-m1 and D2A1-m2 (c) cells were pre-treated with 10 μM FADS2i or FADS1/2i for 4 hours, administrated ON with 0.5 μM erastin and subjected to cell viability assay. One-way ANOVA; Dunnett corrected. (d) Lipid peroxidation level was evaluated by measuring the fluorescence intensity of BODIPY^581/591^-C11 using flow cytometry analysis in the indolent (4T07) and the highly (4T1) metastatic cells pre-treated with 10 μM FADS2i or FADS1/2i for 4 hours, administrated ON with 1 μM RSL3 for 2 hours. Data represent means ± SEMs. One-way ANOVA; Dunnett corrected. (e) Lipid peroxidation level was evaluated by measuring the fluorescence intensity of BODIPY^581/591^-C11 using confocal microscopy in the highly aggressive 4T1 cells pre-treated with 10 μM FADS1/2i for 4 hours, administrated with 5 μM erastin for 2 hours. (Representative confocal images are shown. Oxidized lipids: green; non-oxidized lipids: red; nuclei: blue, DAPI; scale bar, 10 μm). Data represent means ± SEMs. One-way ANOVA; Dunnett corrected. (f) Lipid peroxidation level was evaluated by measuring the fluorescence intensity of BODIPY^581/591^-C11 using cytofluorimetric analysis in D2A1-m1 and D2A1-m2 cells pre-treated with 10 μM FADS2i or FADS1/2i for 4 hours, administrated ON with 1 μM RSL3 2 hours. Data represent means ± SEMs. One-way ANOVA; Dunnett corrected. (g) Human breast cancer cells were treated with 0.5 μM RSL3 for 24 hours and subjected to cell viability assay. (h) MDA-MD-231 cells were pre-treated with 15 μM Fer-1 for 4 hours and then administrated ON with 0.5 μM RSL3 or 0.5 μM erastin. After 24 hours cells were subjected to cell viability assay. One-way ANOVA; Dunnett corrected. (i) Total protein lysates from human TNBC cells were subjected to WB analysis with the antibodies indicated. (j) Human MDA-MB-231 breast cancer cells were pre-treated with 10 μM FADS2i or FADS1/2i for 4 hours, administrated ON with 0.5 μM RSL3 and subjected to cell viability assay. One-way ANOVA; Dunnett corrected. (k) The highly metastatic MDA-MB-231 cells were pre-treated with 10 μM FADS2i or FADS1/2i for 4 hours, administrated ON with 2.5 μM erastin, and subjected to cell viability assay. One-way ANOVA; Dunnett corrected. (l-n) Kaplan-Meier analysis of relapse free survival (RFS) and overall survival (OS) of a curated cohort of TNBC patients divided into high and low as described in Material and Methods section for FADS1 (l) and FADS2 (n) expression. HR and log-rank Mantel-Cox p values are shown. Student’s t test; One-way ANOVA; Each dot represents a biological replicate. * *P*<0.05; ** *P*<0.01; *** *P*<0.001; **** *P*<0.0001.

